# Biological Insights Into Muscular Strength: Genetic Findings in the UK Biobank

**DOI:** 10.1101/201020

**Authors:** Emmi Tikkanen, Stefan Gustafsson, David Amar, Anna Shcherbina, Daryl Waggott, Euan A. Ashley, Erik Ingelsson

**Affiliations:** Division of Cardiovascular Medicine, Department of Medicine, Stanford University School of Medicine, CA; Department of Medical Sciences, Molecular Epidemiology and Science for Life Laboratory, Uppsala University, Sweden

**Author notes:** <u>Address for Correspondence:</u> Erik Ingelsson, MD, PhD, FAHA, 300 Pasteur Dr, mail code: 5773; Stanford, CA 94305; USA.

**Keywords:** Genome-wide association, genetics, grip strength, fitness

## Abstract

**Background:** Hand grip strength, a simple indicator of muscular strength, has been associated with a range of health conditions, including fractures, disability, cardiovascular disease and premature death risk. Twin studies have suggested a high (50-60%) heritability, but genetic determinants are largely unknown.

**Aims:** In this study, our aim was to study genetic variation associated with muscular strength in a large sample of 334,925 individuals of European descent from the UK Biobank, and to evaluate shared genetic aetiology with and causal effects of grip strength on physical and cognitive health.

**Methods and Results:** In our discovery analysis of 223,315 individuals, we identified 101 loci associated with grip strength at genome-wide significance (*P*<5×10^−8^). Of these, 64 were associated (*P*<0.01 and consistent direction) also in the replication dataset (N=111,610). Many of the lead SNPs were located in or near genes known to have a function in developmental disorders (*FTO*, *SLC39A8*, *TFAP2B*, *TGFA*, *CELF1*, *TCF4*, *BDNF*, *FOXP1*, *KIF1B*, *ANTXR2*), and one of the most significant genes based on a gene-based analysis (*ATP2A1*) encodes SERCA1, the critical enzyme in calcium uptake to the sarcoplasmic reticulum, which plays a major role in muscle contraction and relaxation. Further, we demonstrated a significant enrichment of gene expression in brain-related transcripts among grip strength associations. Finally, we observed inverse genetic correlations of grip strength with cardiometabolic traits, and positive correlation with parents’ age of death and education; and showed that grip strength was causally related to fitness, physical activity and other indicators of frailty, including cognitive performance scores.

**Conclusions:** In our study of over 330,000 individuals from the general population, the genetic findings for hand grip strength suggest an important role of the central nervous system in strength performance. Further, our results indicate that maintaining good muscular strength is important for physical and cognitive performance and healthy aging.

## Introduction

Hand grip strength is a simple and non-invasive measurement of general muscular strength and it has been shown to predict disability in older adults, fracture risk, nutritional status, cardiovascular disease events and all-cause mortality^1–3^. Several behavioral and environmental factors, such as physical activity and nutrition, affect the variability of grip strength, but family studies have suggested that genetic factors also have a significant role^4,5^. The identification of genetic variants affecting grip strength variability could help in the understanding of biological mechanisms of muscular fitness, as well as lend biological insights to physical functioning late in life and healthy aging.

So far, two genome-wide association studies (GWAS) of maximal grip strength have been conducted^6,7^. The largest study, also conducted in the UK Biobank, included 195,180 individuals and identified 16 loci associated with grip strength. In this study, we conducted traditional and gene-based GWAS for 334,925 individuals from the UK Biobank to discover novel loci for relative grip strength^8,9^, and evaluated shared genetic aetiology and causal effects of grip strength on physical and cognitive health.

## Results

### Genetic associations for grip strength in biologically relevant loci

In our discovery GWAS (random 2/3 sample from eligible individuals; N=223,315) adjusted for age, sex, genotype array, and 10 principal components, we identified 101 genome-wide significant associations for grip strength (**Figure 1 A**). Four variants were independent loci from the same regions, identified through conditional analysis (rs62106258 near *LINC01874*, rs78648104 in *TFAP2D*, rs800895 in *TRPS1* and rs10871777 near *ENSG00000267620*). Out of 101 variants, 64 were associated (P<0.01 and consistent direction) also in the replication dataset (remaining 1/3 of eligible individuals; N=111,610; **Table 1**). Most of the 64 replicating SNPs were located in introns (48%) or in intergenic regions (22%) while only 8% were located in exons. The two loci with most significant associations were located in chromosome 16, in the intron of *FTO* (rs1421085, β=−0.004, P=5.4×10^−38^ and β=−0.004, P=4.9×10^−22^ in discovery and replication samples, respectively) and *ATXN2L* (rs12928404, β=−0.003, P=1.0×10^−25^ and β=−0.002, P= 1.1×10^−7^). The *FTO* locus has been previously shown to be associated with obesity^10,11^ and lipids^12^, among other metabolic traits; and according to the OMIM database, mutations in this gene can also cause growth retardation, developmental delay, and facial dysmorphism. The other locus in chromosome 16 has been reported to be associated with intelligence in a recent large study^13^, and our lead variant has a high probability of being regulatory (Regulome score^14^=1b). This is also in close vicinity of *ATP2A1*, a gene involved in muscular contraction and relaxation and a causal gene for a muscle disorder called Brody disease, which is characterized by muscle cramping after exercise. A gene-based analysis of the discovery sample identified *ATP2A1* as the most significant gene for grip strength (P=3.9×10^−31^, **Figure 1 B**).

**Figure 1.**
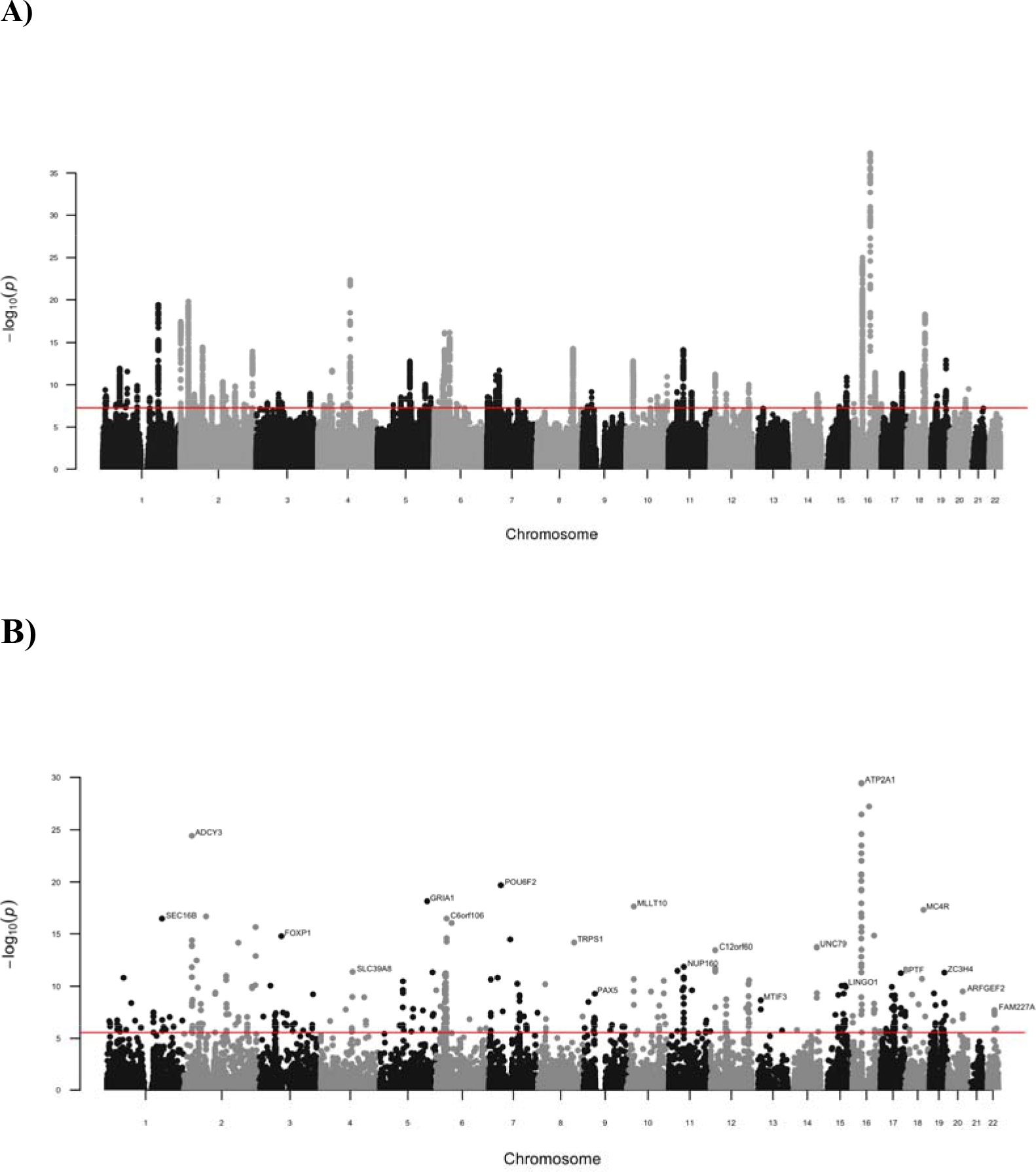
A) Manhattan plot for genetic associations for grip strength in discovery sample. B) Gene-based manhattan plot for genetic associations for grip strength in discovery sample. (The most significant gene for each chromosome labeled)

**Table 1.**
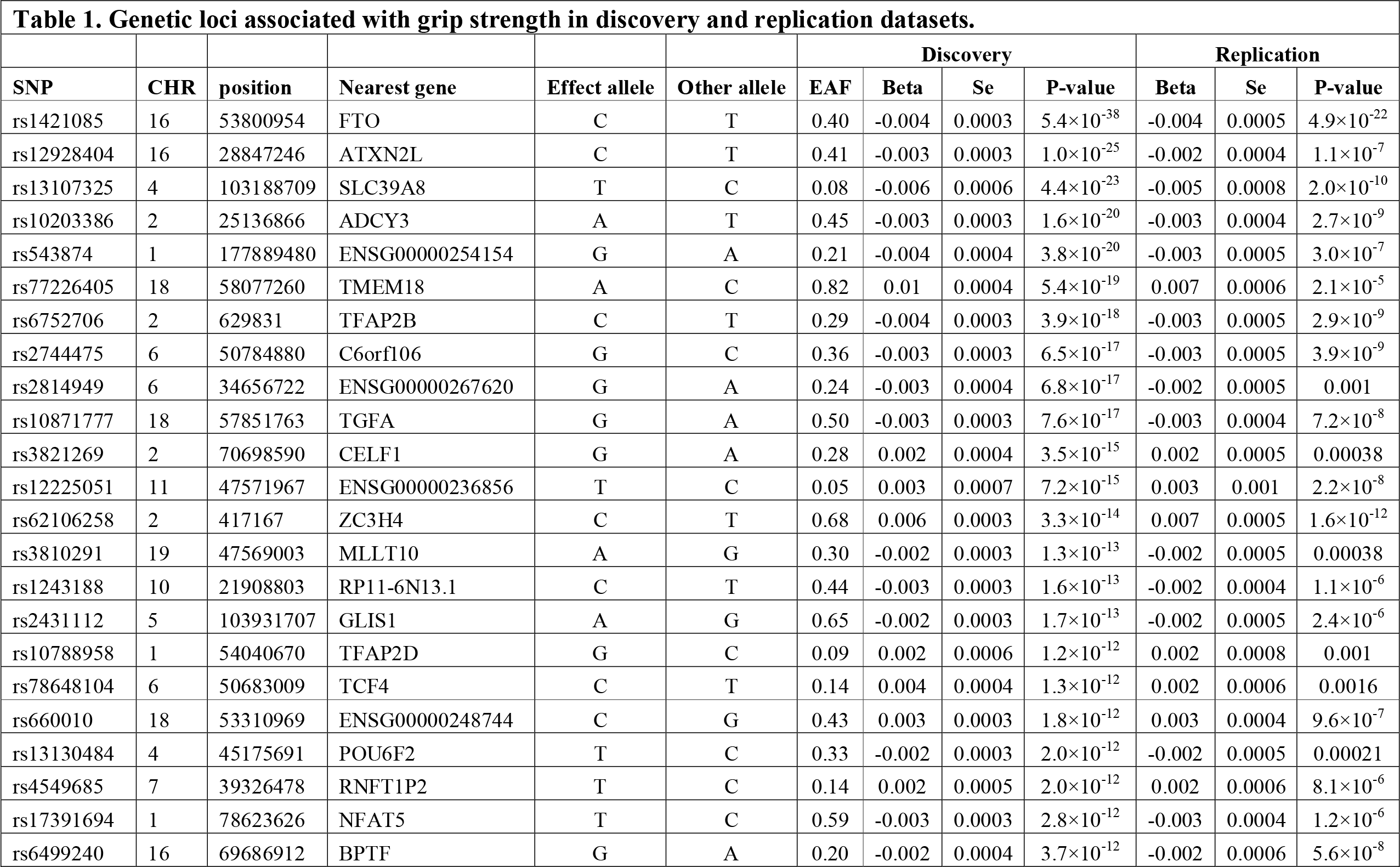
Genetic loci associated with grip strength in discovery and replication datasets.

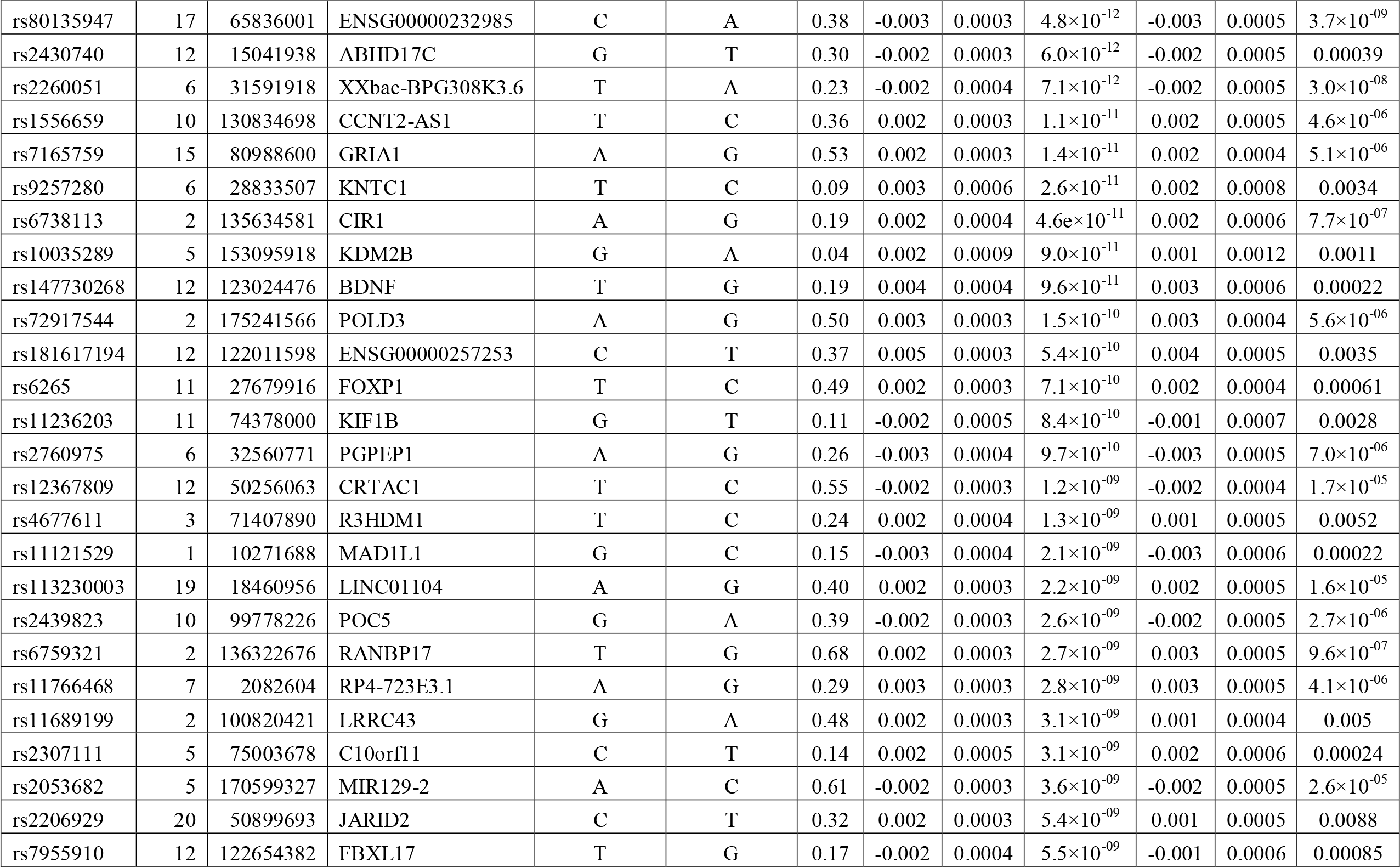

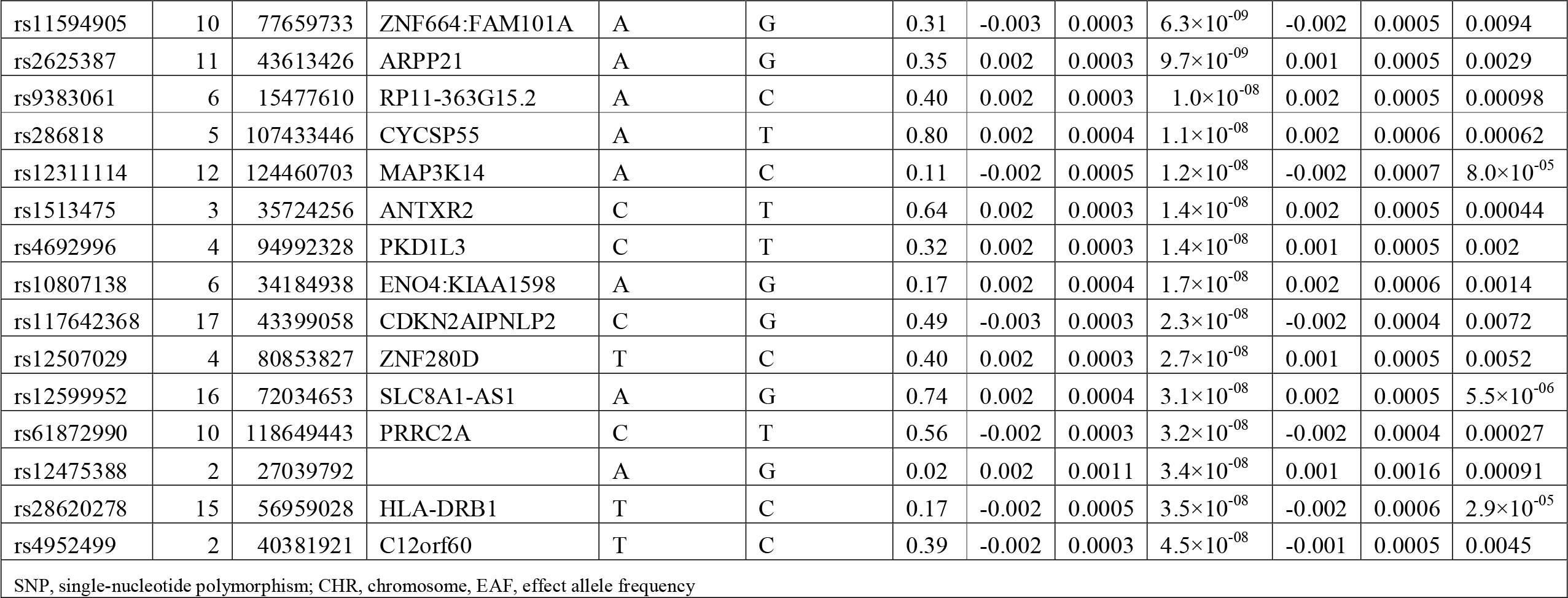

A nonsynonymous SNP in the exon of *SLC39A8* (rs13107325, β=−0.006, P=4.4×10^−23^ and β=−0.005, P= 2.0×10^−10^), has been identified as a susceptibility variant for schizophrenia^15^ and metabolic traits^10,16^. This variant had a CADD score^17^ of 34.00, indicating deleterious effect, and mutations in *SLC39A8* are also known to cause a severe congenital disorder of glycosylation, characterized by delayed psychomotor development apparent from infancy, hypotonia, short stature, seizures, visual impairment, and cerebellar atrophy. Many other genes located in the loci we identified to be associated with grip strength also play roles in different developmental disorders (**Table S1**). These include patent ductus arteriosus, a form of congenital heart defect (*TFAP2B*), cleft lip (*TGFA*), myotonic dystrophy type 1, which is the most prevalent adult onset muscular dystrophy (*CELF1*), Pitt-Hopkins syndrome, which is characterized by intellectual disability, distinctive facial features, poor muscular development and abnormal breathing (*TCF4*), mental retardation with language impairment and with or without autistic features (*FOXP1*), Charcot-Marie-Tooth disease, characterized by distal limb muscle weakness and atrophy due to peripheral neuropathy (*KIF1B*), hyaline fibromatosis syndrome, characterized by abnormal growth of hyalinized fibrous tissue (*ANTXR2*), and congenital central hypoventilation syndrome (*BDNF*). Our top SNP in *BDNF*(rs6265, β=0.002, P=7.1×10^−10^), is located in the exon of *BDNF* and has a high CADD score (CADD=24.1). This is an interesting gene as it encodes brain-derived neurotrophic factor, an important growth factor promoting neurogenesis. BDNF concentration are increased in response to exercise and decreased in neurodegenerative diseases^18^.

To identify candidate genes regulated by the most associated grip strength variants, we used the Genotype-Tissue Expression (GTEx) database^19^ for eQTL analyses. In 25 loci, we found evidence of at least one significant eQTL (FDR≤0.05, **Table S1**). The largest number of significant eQTLs was associated with gene expression in nerve (tibial), artery (tibial), and skin (sun-exposed lower leg). Seven variants had the most significant eQTL for the genes they we located in (*ADCY3*, *TGFA*, *BDNF*, *KIF1B*, *LRRC43*, *ARPP21*, *KIAA1598*).

Interestingly, some variants regulated the expression of other genes having a role in developmental abnormalities. The most significant eQTL was one of the lead variants, rs12928404 in *ATXN2L*, which had a significant association with the expression of adjacent *TUFM* gene in several tissues (lowest P=5.2×10^−69^, FDR=0.0003 for whole blood). Mutations in *TUFM* have previously been shown to cause combined oxidative phosphorylation deficiency 4, a syndrome consisting of intrauterine growth retardation, developmental regression, hypotonia and respiratory failure. Another interesting eQTL was rs6759321, which regulated *DARS* expression in thyroid (FDR=0.0002). *DARS* is a causal gene for another disorder including delayed motor development, mental retardation, among other features (“hypomyelination with brainstem and spinal cord involvement and leg spasticity”). Further, rs12599952 regulated the expression of *DHODH* in several tissues (lowest P=1.6×10^−11^, FDR=0.0003 for tibial artery), a gene responsible for Miller syndrome including severe micrognathia and several other developmental abnormalities.

### Genetic risk score analysis

We then calculated a genetic risk score (GRS) as a weighted sum of the 101 grip strength variants identified in the discovery dataset and estimated its associations with the measures of fitness, general health and indicators of frailty^20^ in the replication dataset. The GRS was significantly associated cardiorespiratory fitness (VO_2_, β=0.20 per SD in GRS, SE=0.02, P=3.1×10^−19^), objective measurement of physical activity (average acceleration measured with a wrist-worn accelerometer, β=0.02, SE=0.006, P=0.003), self-reported good or excellent overall health (β=0.07, SE=0.007, P=1.3×10^−22^), and fluid intelligence score (β=0.01, SE=0.005, P=0.005). The significant inverse associations were observed with slow walking speed (β=-0.11, SE=0.01, P=1.1×10^−20^), frequent feelings of tiredness / lethargy in last 2 weeks (β=−0. 03, SE=0.006, P=1.4×10^−5^), falls during the last year (β=−0.04, SE=0.008, P=8.5×10^−8^), weight loss during the last year (β=−0.04, SE=0.008, P=8.5×10^−8^), and reaction time (β=−0.01, SE=0.003, P=1.6×10^−5^). All association remained significant after multiple testing correction (alpha threshold = 0.05/9).

To evaluate the causality of grip strength on these outcomes, we applied Mendelian Randomization (MR) analysis with two-stage regression. This analysis supported causal effects for all tested outcomes (**Table 2**). Weak instrument test was rejected for all models, indicating that the GRS was powerful enough to detect the causal effect. Durbin-Wu-Hausman test indicated significant endogeneity for cardiorespiratory fitness and weight loss, suggesting the presence of some unmeasured confounding.

**Table 2.** Causal effects of grip strength on frailty indices.

| **Table 2. Causal effects of grip strength on frailty indices.** |  |  |  |  |  |
| --- | --- | --- | --- | --- | --- |
| **Trait** | **Beta\*** | **Se** | **P** | **Weak instruments P-value** | **Durbin-Wu-Hausman** |
| Physical activity (accelerometer) | 0.230 | 0.073 | 0.0017 | <0.001 | 0.95 |
| Cardiorespiratory fitness (VO2) | 2.569 | 0.282 | $8.3 \times 10^{-20}$ | <0.001 | <0.001 |
| General Health | 0.150 | 0.015 | $1.1 \times 10^{-23}$ | <0.001 | 0.099 |
| Slow walking speed | -0.087 | 0.009 | $1.6 \times 10^{-21}$ | <0.001 | 0.07 |
| Falls | -0.076 | 0.014 | $4.7 \times 10^{-08}$ | <0.001 | 0.13 |
| Weight loss | -0.103 | 0.013 | $2.3 \times 10^{-15}$ | <0.001 | <0.001 |
| Tiredness | -0.077 | 0.017 | $9.9 \times 10^{-06}$ | <0.001 | 0.91 |
| Fluid intelligence score | 0.178 | 0.063 | 0.0049 | <0.001 | 0.078 |
| Reaction time | -0.139 | 0.033 | $2.7 \times 10^{-05}$ | <0.001 | 0.57 |
| * Per SD change in grip strength. VO2: net oxygen consumption |  |  |  |  |  |

### Meta-analysis

To maximize power for pathway analyses and to suggest additional associations, we also performed a meta-analysis of the discovery and replication samples. This analysis revealed 139 independent loci reaching genome-wide significance for grip strength (**Table S2**). These variants combined explained 1.1% of the grip strength variance in the UK Biobank (which could constitute an overestimation due to Winner’s curse, as the variants were discovered in this sample). Based on LD-score regression of meta-analyzed results, the genome-wide “chip” heritability of grip strength was 0.13 (SE◻ =◻0.004). We observed significant inflation of P-values (λ_GC_=1.55), but the LD-score regression intercept of 0.9 suggested that this was due to polygenicity, rather than population stratification.

Using the full distribution of SNP P-values from the meta-analysis, we observed a significant positive relationship between genes highly expressed in brain and genetic associations for grip strength (**Figure 2**). When using only the prioritized genes (based on positional mapping) in meta-analysis as input genes, we observed the most significant enrichment of differentially expressed genes in muscle (P=0.003 for up and down regulated, adjusted P=0.10). Further, the proportion of overlapping genes in GO biological processes gene sets was highest for “go regulation of skeletal muscle contraction” (overlapping genes *GSTM2*, *CASQ1*, *ATP2A1*, P=6.3×10^−5^, adjusted P=0.009), which is described as any process that modulates the frequency, rate or extent of skeletal muscle contraction.

**Figure 2.**
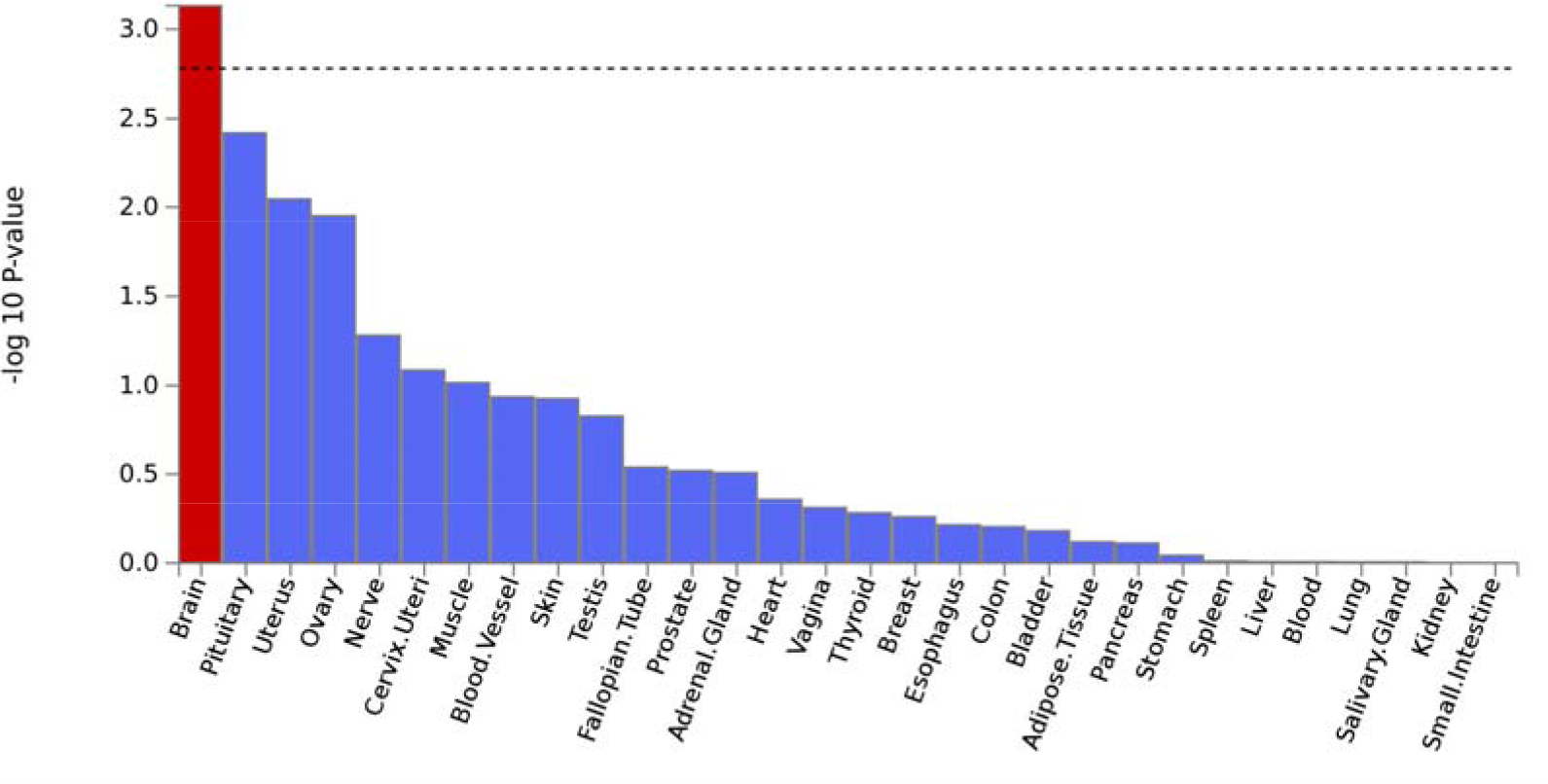
Tissue-expression of grip strength loci.

### Genetic correlations

To evaluate genome-wide heritability 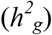, partitioned heritability by functional categories, and genetic correlation between other traits, we applied LD-score regression^21–24^ on the results from our meta-analysis. In line with 17 complex diseases and traits analyzed by Finucane et al.^24^, we identified strong enrichment of grip strength loci in conserved regions (proportion of heritability / proportion of SNPs = 15.6, P=5.0×10^−20^, FDR=2.6×10^−18^). When testing for genetic correlations between grip strength and all traits in LD Hub^23^, we observed significant genetic correlations with 78 traits (**Figure 3**), of which most of them were cardiometabolic traits. The strongest negative correlations were observed with obesity measures and leptin. Interestingly, strong negative correlations were also detected with attention deficit hyperactivity disorder and depressive symptoms. The strongest positive correlations were observed with parent’s age at death, high-density lipoprotein, years of schooling and forced vital capacity.

**Figure 3.**
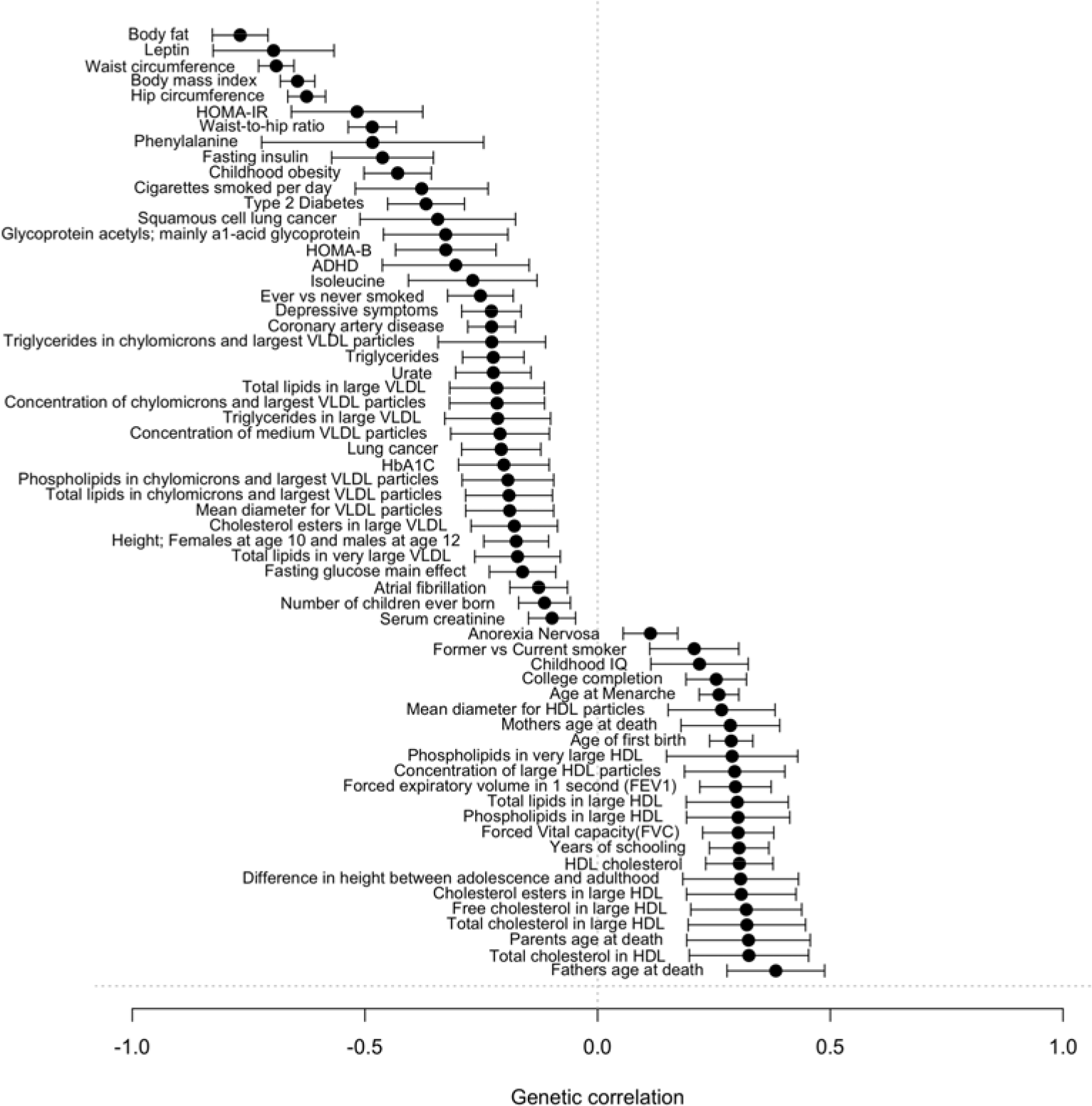
Genetic correlation between grip strength and other traits.

## Discussion

In this largest GWAS to date for grip strength, we report 64 loci robustly associated with grip strength. Many of the lead SNPs were located in or near genes known to have a function in developmental disorders, and one of the most significant genes based on gene-based analysis (*ATP2A1*) encodes SERCA1, the critical enzyme in calcium uptake to the sarcoplasmic reticulum, which plays a major role in muscle contraction and relaxation. The largest number of significant eQTLs was observed for tibial nerve and several genes regulated by our top SNPs play a role in neuro-developmental abnormalities or brain function. Further, grip strength was causally related to fitness, physical activity and other indicators of frailty, including cognitive performance scores. In our meta-analysis of discovery and replication samples, the number of significant loci reached 139 independent variants, and we showed that genes in these loci have a significant enrichment of gene expression in brain. Finally, we observed inverse genetic correlations with cardiometabolic traits, but also attention deficit hyperactivity disorder and depressive symptoms; and positive correlation with parents’ age of death and education.

Our results allow us to draw several conclusions. First, it is well known that muscular strength depends not only on the quantity of the involved muscles, but also on the ability of the nervous system to appropriately recruit the muscle cells^25^. Thus, this is concordant with our results showing the brain as the most important organ regulating muscular strength. Second, the underlying biology of grip strength points to genes with a known role in muscle and brain function, and neuro-developmental disorders, many of them which are characterized by disturbances in motor development and intellectual disability. Further, our causal and genetic correlation analyses suggest shared genetic aetiology between muscular strength and cognitive performance, which is concordant with observational and experimental findings about the beneficial effects of exercise training on brain health^18^. The underlying biological mechanisms could relate to the stimulating effects of exercise in the brain, promoting neuroplasticity. This hypothesis also gets support in an evolutionary context^26^. Finally, the existing literature shows that higher grip strength is associated with lower risk of all-cause and cardiovascular mortality^1^, which is consistent with our findings of high positive genetic correlation with parents’ age of death, negative correlation with several cardiometabolic traits and causal effects of grip strength on frailty. These findings suggest that maintaining good strength has a key role in physiological functioning and healthy aging.

The main strengths of this study include a very large sample size, which enabled us to detect a much larger number of genetic variants compared to previous studies^6,7^. We also used relative grip strength as our phenotype, which is less confounded by body size and more suitable to reflect general muscular fitness than maximal grip strength. Our study also has some limitations. First, the response rate in UK Biobank was low (5.5%), and the generalizability of results is unknown^27^. Second, even though grip strength is a commonly used proxy for muscular fitness, it captures mainly upper body strength, especially when measured in sitting position. However, it is highly correlated with knee extension muscle strength (r= 0.77 to 0.81)^28^; and due its feasibility, it is convenient proxy for muscular strength in large samples, which are required in genetic association studies. Finally, our GWAS were conducted in unrelated European samples only to avoid population stratification. Thus, the generalizability to other ethnicities is unknown.

In conclusion, we identified a large number of genetic loci associated with hand grip strength providing insights of biological processes involved in individual variation in muscular strength and providing clues for discovery of new treatments for muscle-related diseases. Our results highlight the central role of nervous system in strength performance and the importance of maintaining muscular strength to prevent age-related physical and cognitive decline.

## Material and Methods

### Study sample

The UK Biobank is a large longitudinal cohort including over 500,000 individuals aged 40-69 years. Participants were enrolled in 22 study centers located in England, Scotland and Wales during 2006-2010. Extensive baseline data on medical history, health behavior, and physical measurements were collected by questionnaires and clinical examination. Participants also provided samples blood, urine and saliva and have agreed to have their future health, including disease events, monitored. The UK Biobank study was approved by the North West Multi-Centre Research Ethics Committee and all participants provided written informed consent to participate in the UK Biobank study. The study protocol is available online^29^.

### Phenotypes

Grip strength was measured in a sitting position using a Jamar J00105 hydraulic hand dynamometer. This measures grip force without movement and adjusts the participant’s hand size. The participants were asked to squeeze the device as hard as they could for three seconds, and the maximum value that was reached during that time was recorded. Both hands were measured in turn (UK Biobank field ID 46 for left and 47 for right hand). Due to its high correlations with body size measurements, relative grip strength (absolute strength corrected for body size) has been suggested to be more accurate measurement of strength. Strength might be higher in obese individuals, but the relative strength (muscle strength per kilogram of body weight) is much lower^8,9,30^. Thus, we calculated relative grip strength as an average of measurements of right and left hand divided by weight (ID 21002)^8,9^. Weight was measured with bioelectrical impedance analysis (BIA) Tanita BC418MA. Standing height was measured using a Seca 202 device for participants standing barefoot. Objective measurement of physical activity was measured with Axivity AX3 wrist-worn triaxial accelerometer^31^. Cardiorespiratory fitness was measured with the cycle ergometry on a stationary bike (eBike, Firmware v1.7). We calculated net oxygen consumption (VO_2_) from individuals’ body weight and maximum workload using the equation VO_2_=7+10.8(workload)/weight^32^. Fluid intelligence score was calculated as a sum of the number of correct answers given to the 13 fluid intelligence questions.

Reaction time was determined as mean time to correctly identify matches in snap reaction speed game. The categorical variables of self-reported general health and frailty indicators were recoded to binary variables; overall health: good or excellent, 1, others, 0; slow walking speed: slow pace, 1, others, 0; feelings of tiredness / lethargy in last 2 weeks was defined: more than half of the days or more frequently, 1, others, 0; falls during the last year: one or more falls, 1, others, 0; and weight loss during the last year, yes – lost weight, 1, others, 0.

### Genotypes

Genotyping was performed with the UK BiLEVE and UK Biobank Axiom arrays (Affymetrix Research Services Laboratory, Santa Clara, California, USA) including 807,411 and 825,927 markers, respectively. Initial quality control (QC) was conducted centrally by the UK Biobank, and has been described in detail by Bycroft et al^33^. In short, poor quality genetic markers were detected based on statistical tests for batch effects, plate effects, departures from Hardy-Weinberg Equilibrium (HWE), sex effects, array effects, and discordance. Poor quality samples were identified based on the metrics of missing rate and heterozygosity. After quality control, the data consisted of 488,377 samples (N=49,950 and 438,427 individuals with the UK BiLEVE and UK Biobank Axiom arrays, respectively) and 805,426 single nucleotide polymorphisms (SNPs), which were the imputed with IMPUTE2 by using both HRC and 1000 Genomes Phase 3 merged with the UK10K haplotype reference panels, so that the HRC was preferred for SNPs present in both panels. In our analysis, we used July 2017 release of the imputed genetic marker data, by excluding genetic markers imputed with the UK10K + 1000 Genomes reference panel due to reported imputation error. We further excluded genetic markers with minor allele count ≤30 and imputation quality <0.8. Thus, the total number of genetic markers included in our analysis was 15,275,733. Further, we included only unrelated individuals with selfreported British descent and European/Caucasian ethnicity based on principal component analysis.

### Association analysis

The discovery GWAS was carried out in the random sample of 223,315 individuals. Analysis was performed with a linear regression by using PLINK^34^ (version 2.0) assuming an additive model for association between phenotypes and genotype dosages. For top loci, we performed a conditional analysis for a region around the lead SNP to identify additional independent variants. For replication, we used the remaining data of 111,610 individuals. Age, sex, genotype array and 10 principal components were used as covariates. Finally, we conducted an inverse-variance weighted fixed-effect meta-analysis with METAL software^35^ to reveal additional genetic variants associated with grip strength.

### Functional annotation

We mapped the genomic positions of all replicated lead SNPs from our discovery analysis and annotated them with the closest gene with ANNOVAR. We used OMIM database to search for known genetic disorders for mapped genes. In addition, we obtained CADD scores^17^, the score of deleteriousness of SNPs predicted by 63 functional annotations, and RegulomeDB scores^14^, representing regulatory functionality of SNPs based on eQTLs and chromatin marks. SNPs were also mapped with tissue-specific gene expression levels by utilizing GTEx database^19^. We searched for significant eQTLs (FDR ≤ 0.05) across all available tissues.

Gene-based association test was computed by MAGMA (v. 1.6)^36^, by mapping SNPs into 18,225 protein-coding genes, and then testing the joint association of all markers in the gene with grip strength with a multiple linear principal components regression. Bonferroni-correction was used to define genome-wide significance (alpha threshold = 0.05/18,225). To evaluate enrichment in tissue-specific gene expression, we tested for association between all genetic associations from meta-analysis and gene expression in a specific tissue types by averaging gene-expression per tissue type. Gene set enrichment analysis was conducted for prioritized genes (based on position) in meta-analysis. Gene sets were obtained from MsigDB (v. 5.2) and multiple test correction was performed for each category.

Analyses were conducted with FUMA platform^37^ using functions SNP2GENE and GENE2FUNC.

### Genetic risk score analyses

We included 101 independent, genome-wide significant SNPs identified in our discovery sample for our GRS analysis. The GRS was calculated as a weighted sum of the grip strength increasing alleles, by using the effect sizes from the discovery analysis as weights. We evaluated the association between the GRS and cardiorespiratory fitness, objective measurement of physical activity, fluid intelligence score and reaction time with linear regression models adjusted for age, sex, genotype array and 10 principal components. The outcome variables were rank transformed to normal distribution. The associations between the GRS and selfreported overall health, slow walking speed, frequent feelings of tiredness / lethargy in last 2, falls during the last year, and weight loss during the last year were analyzed with logistic regression models adjusted for age, sex, genotype array and principal components.

We then performed causal analysis with two-stage regression. In the first stage, a conventional regression was used to assess the association between the GRS and observed grip strength (adjusting for the same covariates as above). Then, the predicted value of the trait from the model was used as an independent variable in the second stage, where the dependent variable was the trait of interest (e.g. physical activity). The beta coefficient from the second regression reflects an unconfounded effect of genetically determined grip strength on the outcome. Endogeneity, the difference between the causal estimate and the observed effect size, was examined by Durbin-Wu-Hausman. The analysis was conducted with the R-package AER.

### LD-score regression

By leveraging genome-wide information from our meta-analysis, we used LD-score regression^21–24^ to estimate heritability of grip strength, to evaluate partitioned heritability by functional categories and to identify genetic correlation with other traits. LD-score regression was conducted using the summary statistics from the metaanalysis of discovery and replication. We used pre-calculated European LD scores and restricted SNPs to those found in HapMap Phase III to ensure good quality imputation. We conducted partitioned heritability analysis using 24 main annotations described by Finucane et al.^24^. Enrichment was defined as the proportion of SNP heritability explained divided by the proportion of SNPs in each functional category. We considered FDR ≤ 0.05 to indicate statistically significant enrichments. Genetic correlation was tested against all traits in LD Hub^23^. To identify significant genetic correlations, we set the P-value threshold based on Bonferroni correction (0.05/231).

## Acknowledgements

This research has been conducted using the UK Biobank Resource under Application Number 13721. The Genotype-Tissue Expression (GTEx) Project was supported by the Common Fund of the Office of the Director of the National Institutes of Health, and by NCI, NHGRI, NHLBI, NIDA, NIMH, and NINDS. The data used for the analyses described in this manuscript were obtained from the GTEX Portal on 08/28/17. Some of the computing for this project was performed on the Sherlock cluster. We would like to thank Stanford University and the Stanford Research Computing Center for providing computational resources and support that contributed to these research results. The research was performed with support from National Institutes of Health (1R01HL135313-01; 1R01DK106236-01A1), Knut and Alice Wallenberg Foundation (2013.0126), Finnish Cultural Foundation, Finnish Foundation for Cardiovascular Research and Emil Aaltonen Foundation.

## Conflicts of interests

Erik Ingelsson is a scientific advisor for Precision Wellness and Olink Proteomics for work unrelated to the present project.

## Authors' contributions

Design of the study: ET, DA, AS, DW, EA, EI; Analysis and interpretation of data: ET, SG; Drafting the manuscript: ET; Critical revision of the manuscript: all authors; Final approval of the version to be published: all authors.

